# *In vivo* control of *Toxoplasma gondii* by zebrafish macrophages

**DOI:** 10.1101/828624

**Authors:** Nagisa Yoshida, Marie-Charlotte Domart, Artur Yakimovich, Maria J. Mazon-Moya, Lucy Collinson, Jason Mercer, Eva-Maria Frickel, Serge Mostowy

## Abstract

*Toxoplasma gondii* is an obligate intracellular parasite capable of invading any nucleated cell. Three main clonal lineages (type I, II, III) exist and murine models have driven the understanding of general and strain-specific immune mechanisms underlying *Toxoplasma* infection. However, murine models are limited for studying parasite-leukocyte interactions *in vivo*, and discrepancies exist between cellular immune responses observed in mouse versus human cells. Here, we develop a zebrafish infection model to study the innate immune response to *Toxoplasma in vivo*. By infecting the zebrafish hindbrain ventricle, and using high-resolution microscopy techniques coupled with computer vision driven automated image analysis, we reveal that *Toxoplasma* invades and replicates inside a parasitophorous vacuole to which type I and III parasites recruit host cell mitochondria. We show that type II and III strains maintain a higher infectious burden than type I strains. To understand how parasites are being cleared *in vivo*, we analyzed *Toxoplasma*-macrophage interactions using time-lapse and correlative light and electron microscopy. Strikingly, macrophages are recruited to the infection site and play a key role in *Toxoplasma* control. These results highlight *in vivo* control of *Toxoplasma* by macrophages, and illuminate the possibility to exploit zebrafish for discoveries within the field of parasite immunity.

## Introduction

*Toxoplasma gondii* is a successful human pathogen that often remains asymptomatic, however complications arise in the immunocompromised and in neonates if infection is contracted during pregnancy (1). *Toxoplasma* exist as invasive rapidly replicating tachyzoites in intermediate hosts (such as rodents and livestock), and convert into bradyzoite cysts in immune privileged sites and long-lived cells (such as the brain and muscle tissue) during chronic infection (2). Once inside the host cell, parasites reside in a non-fusogenic parasitophorous vacuole (PV) where *Toxoplasma* asexually replicates (3). Egress leads to dissemination into neighboring tissues, culminating in systemic infection. Predation of intermediate hosts by the definitive feline host completes the *Toxoplasma* life cycle. Control of infection by the host immune response is thus critical for both host survival and for continued parasite transmission. As a result of its well-understood life cycle, *Toxoplasma* has emerged as a valuable model organism to understand the balance of pathogen survival and innate cellular immune control.

Three clonal lineages of *Toxoplasma* dominate across Europe and South America, namely the type I, II and III strains (4). These three closely-related *Toxoplasma* strains have been characterized by the severity of infections they cause in murine models (5). Infection with type I parasites causes acute mouse mortality, whereas infection with type II and type III parasites progress towards chronic infection (6, 7). In humans, it is thought that type II strains predominate in Europe, yet strain-dependent differences in pathogenesis and host responses are poorly understood (8, 9).

Innate immune mechanisms against *Toxoplasma* infection have been studied *in vitro* using both murine and human cell lines and *in vivo* using mice. *In vivo* studies have shown monocytes and neutrophils are recruited to the intestine upon oral infection, and are the major cell types infected with *Toxoplasma* both *in vivo* and *ex vivo* in human peripheral blood (10–13). The importance of neutrophils in parasite control *in vivo* is not fully understood, yet neutrophil-specific depletion studies have suggested a minor protective role against *Toxoplasma* (14, 15). In contrast, inflammatory monocytes are the first responders to infection and are crucial for controlling acute *Toxoplasma* infection (16–18). Pioneering work identified the ability of macrophages to kill *Toxoplasma* (19, 20), by employing both IFN-γ-dependent and -independent mechanisms to control intracellular parasite replication (21–23).

While the mouse is a natural intermediate host and remains an important model to understand *Toxoplasma* pathogenesis, differences are emerging between the mouse and human in mechanisms of parasite control (5, 24–28). Therefore, to complement *in vivo* murine studies, a novel animal model can benefit analysis of *Toxoplasma* control on a cellular and molecular level. Zebrafish are a well-established model for studying infection and immunity (29–32). Coupled with their optical accessibility during early development, zebrafish larvae are highly suited for non-invasive study of *Toxoplasma* infection and host response in real-time *in vivo* (31, 32). Here we develop a zebrafish infection model to study strain-dependent infectivity and leukocyte response to *Toxoplasma infection*. We discover type II (Pru) and III (CEP) parasites maintain a higher infectious burden than type I (RH) parasites. We show macrophages are crucial in the clearance of viable parasites. Our zebrafish infection model can be used as a novel platform to enable unprecedented discoveries in strain-dependent parasite immunity.

## Results

### Intracellular *Toxoplasma* replicate in the zebrafish hindbrain ventricle

To develop a *Toxoplasma*-zebrafish infection model, we tested if tachyzoites could replicate in zebrafish larvae. We first used *Toxoplasma* type I (RH) strain, since it is known to grow faster *in vitro* and survive longer extracellularly than type II (Pru) and type III (CEP) strains (6, 33, 34). We injected zebrafish larvae 3 days post-fertilization (dpf) in the hindbrain ventricle (HBV) with ~5×10^3^ type I strain tachyzoites expressing GFP and followed infection for 24 hours at 33°C (Sup. Fig. 1A). We observed parasite replication *in vivo* using time-lapse widefield fluorescent microscopy (Sup. Fig. 1B, Movie 1). Consistent with this, confocal microscopy shows the percentage of vacuoles containing two or more tachyzoites significantly increasing with time (Fig. 1A and B). To identify the intracellular location of replicating parasites, infected larvae were fixed and stained for granule antigen 2 (GRA2), a dense granule protein that accumulates in the PV lumen (35). Here, GRA2 accumulates around single and replicating parasites, highlighting PV formation *in vivo* (Fig. 1C).

**Fig. 1.**
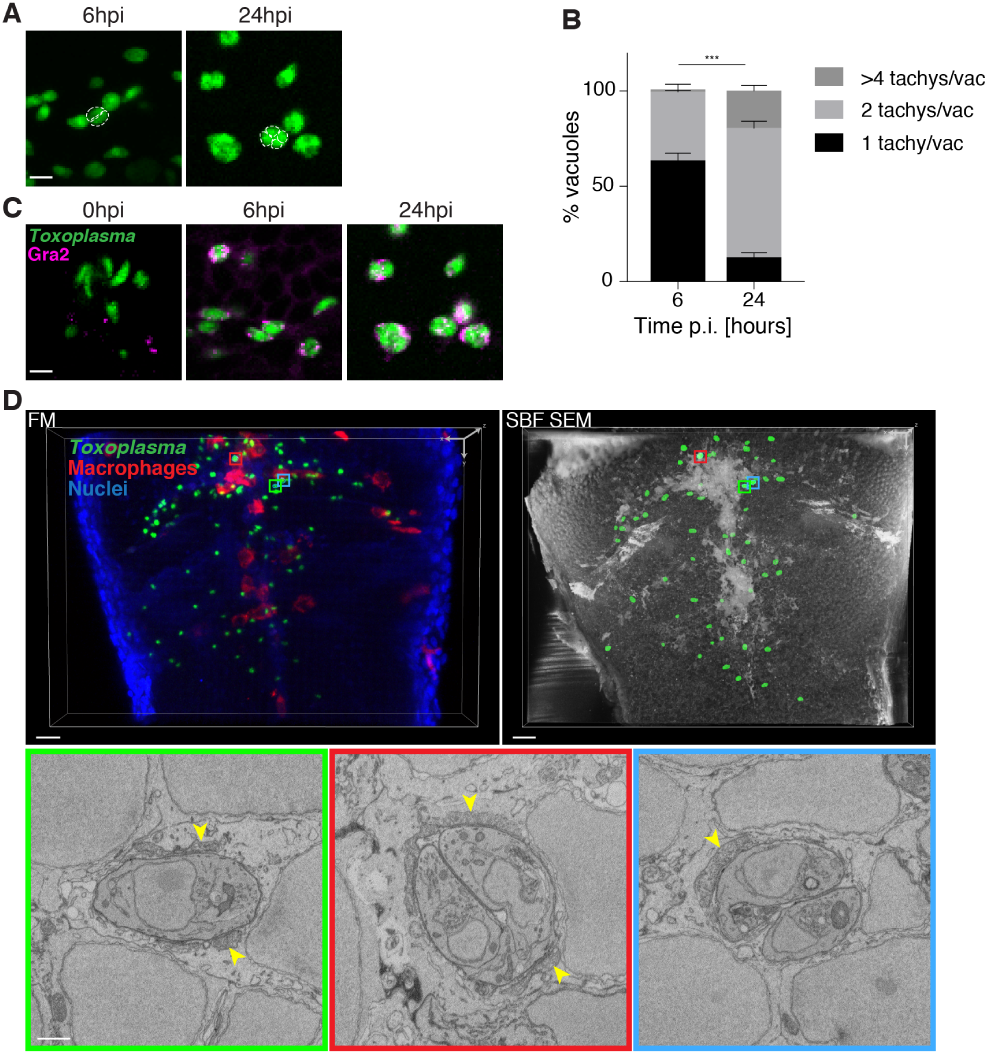
*Toxoplasma gondii* tachyzoites are intracellular and replicate in zebrafish. (A) Representative images from confocal imaging of fixed larvae infected in the HBV with type I *Toxoplasma*-GFP at 6 and 24 hpi. Individual tachyzoites indicated by dashed white outlines. Scale bar, 5 µm. (B) Pixel volume quantification of individual GFP-positive punctae at 6 and 24 hpi. Presented as percentage of total vacuoles counted in the HBV that are 1 tachyzoite/vacuole (<50 pix_3_), 2 tachyzoites/vacuole (50<100 pix_3_) or >4 tachyzoites/vacuole (>100 pix_3_). Pooled data from 3 independent experiments with at least 3 larvae per time point. Significance calculated using 2-way ANOVA, ***, p0.001. Mean ± SEM shown. (C) Representative confocal images of larvae infected in the HBV with type I *Toxoplasma*-GFP (green), fixed at 0, 6 and 24 hpi and labelled with *α*-GRA2 (magenta). Scale bar, 5µm. (D) CLEM of tachyzoites in the HBV of transgenic mpeg1:G/U:mCherry larvae harbouring red macrophages infected with type I *Toxoplasma*-GFP (green) at 6 hpi. 3D reconstructions of 40 confocal z-stacks of a full vibratome section (FM; fluorescence microscopy, top left panel) and of 354 inverted consecutive 50 nm SBF SEM slices of a segment of it (top right panel). A middle slice of each of the *Toxoplasma* visible in the SBF SEM dataset was manually segmented (green, top right panel) to aid correlation. Regions of interest showing the localization of the high-resolution SBF SEM images (lower panels) are denoted with colour boxes. Single (left, green box), replicating (middle, red box) and doublet (right, blue box) tachyzoites in zebrafish host cells were observed. See also Movie 2. Showing three representative images out of a total of 36 *Toxoplasma* in zebrafish cells (see Sup. Fig. 1C) images in their whole volume to accurately determine their stage. Host mitochondrial recruitment to the parasitophorous vacuole indicated by yellow arrowheads. Scale bar, 10 µm (top panels) and 1 µm (lower panels).

To investigate parasite morphology and location at 6hpi, 3D correlative light and electron microscopy (CLEM) was performed on the HBV of infected zebrafish. In this case, we observe that parasites are inside zebrafish cells and display host mitochondrial association (Fig. 1D, other text Sup. Fig. 1C, Movie 2), a hallmark of intracellular type I parasites previously observed in mouse and human cells (36). Moreover, tachyzoites can be observed as singlets, replicating doublets (thus joined together) or fully replicated doublets (with a distinct membrane around each tachyzoite) (Fig. 1D, Sup. Fig. 1C, Movie 2). Collectively, these results show type I *Toxoplasma* tachyzoites can invade zebrafish cells and replicate *in vivo*.

### Type II and III parasites are more efficient than type I parasites at establishing infection *in vivo*

To determine if parasite strain can affect parasite burden and host response in our zebrafish model, we infected larvae with ~5×10^3^ type I, type II or type III *Toxoplasma*-GFP (Sup. Fig. 2A). In all cases, infected larvae show 100% survival and no adverse effects up to 48hpi (Sup. Fig. 2B). Analysis by fluorescent stereomicroscopy showed, from initial parasite input, parasite burden is reduced ~95% by 6hpi, suggesting ~5% of parasites successfully invade zebrafish cells and establish infection. To quantify parasite burden in a high-throughput manner we optimized an automated quantification pipeline using ZedMate (37) for the different strain types at 6 and 24hpi. Strikingly, type II and III parasite burden is ~3x higher than type I parasite burden at 6hpi (Fig. 2A and B, Sup. Fig. 2C). However, once established at 6hpi, all 3 strain types persist equally and decrease by ~20% between 6 and 24hpi.

**Fig. 2.**
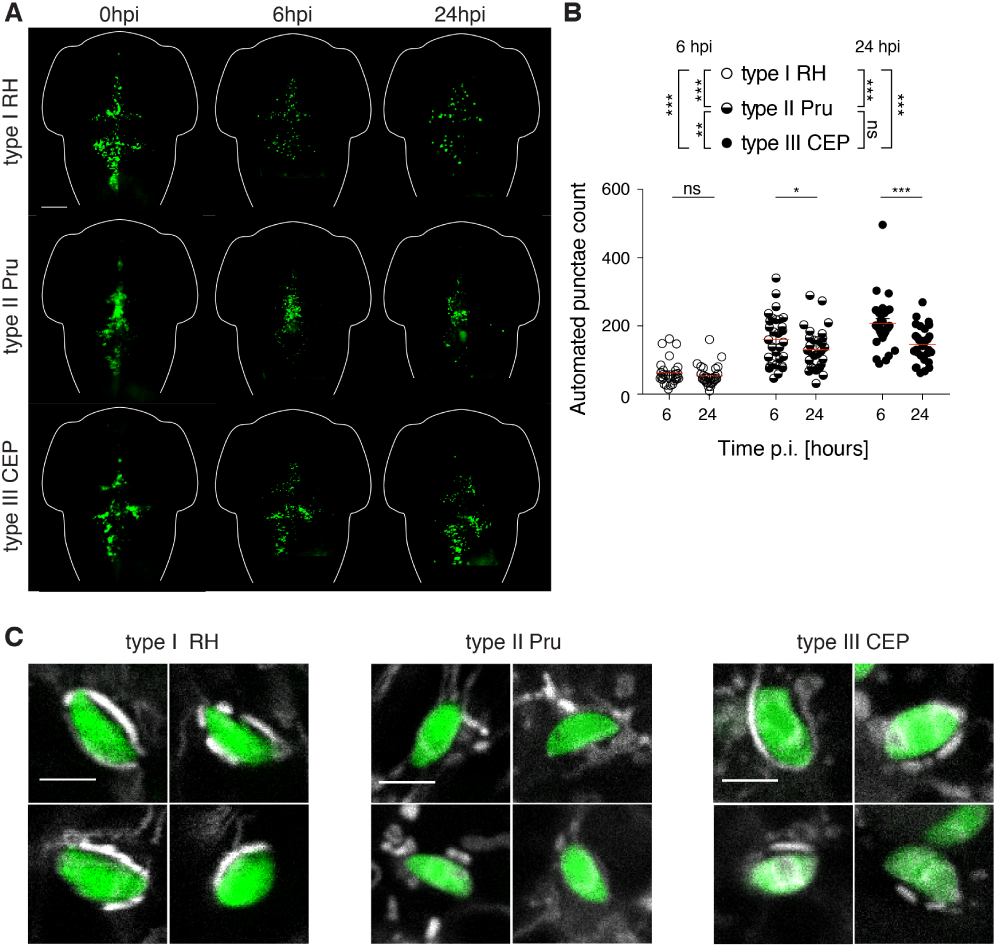
Non-lethal zebrafish larvae model of acute *Toxoplasma gondii* infection. (A) Representative images of larvae infected in the HBV with type I (top panels), type II (middle panels) or type III (bottom panels) of *Toxoplasma* (green). Individual larvae were imaged and monitored at 0, 6 and 24 hpi (from left to right) by fluorescent stereomicroscopy. Scale bar, 100 µm. (B) Automated enumeration of GFP-positive punctae at 6 and 24 hpi of larvae infected with type I (open circle), type II (semi-closed circle) or type III (closed circle of *Toxoplasma* tachyzoites. Automated counts were supported by manual quantifications (Sup. Fig. 2C). Mean ± SEM shown. Pooled data from at least 3 independent experiments with at least 5 larvae per condition per experiment. Significance calculated using 2-way ANOVA, ns, p>0.05, *, p *≤* 0.05, ***, p *≤* 0.001. (C) Representative confocal images of larvae infected in the HBV with type I (left panels), type II (middle panels) or type III (right panels) of *Toxoplasma* (green) and stained for mitochondria (white) 6 hpi. Scale bar, 5 µm.

To test if host mitochondrial association is observed across the three strain types, we stained host mitochondria in the HBV of infected zebrafish larvae. In agreement with *in vitro* observations (36), both type I and type III parasites (and not type II parasites) show clear host mitochondrial association (Fig. 2C). These results demonstrate that strain type-dependent host mitochondrial association characteristics are conserved in zebrafish *in vivo*.

### Macrophage and neutrophil response to parasite infection *in vivo*

To analyze *Toxoplasma*-macrophage interactions over time, 3dpf transgenic larvae possessing red macrophages Tg(*mpeg1:Gal4-FF*)^*gl25*^/Tg(*UAS-E1b:nfsB.mCherry*)^*c264*^ (herein referred to as *mpeg1:G/U:mCherry*), were infected with type I, II or III *Toxoplasma*-GFP, and macrophage recruitment was quantified by fluorescent stereomicroscopy. As compared to mock injection, the number of macrophages recruited to the infection site is significantly increased (~1.5 fold) for all three strain types at both 6 and 24hpi (Fig. 3A).

**Fig. 3.**
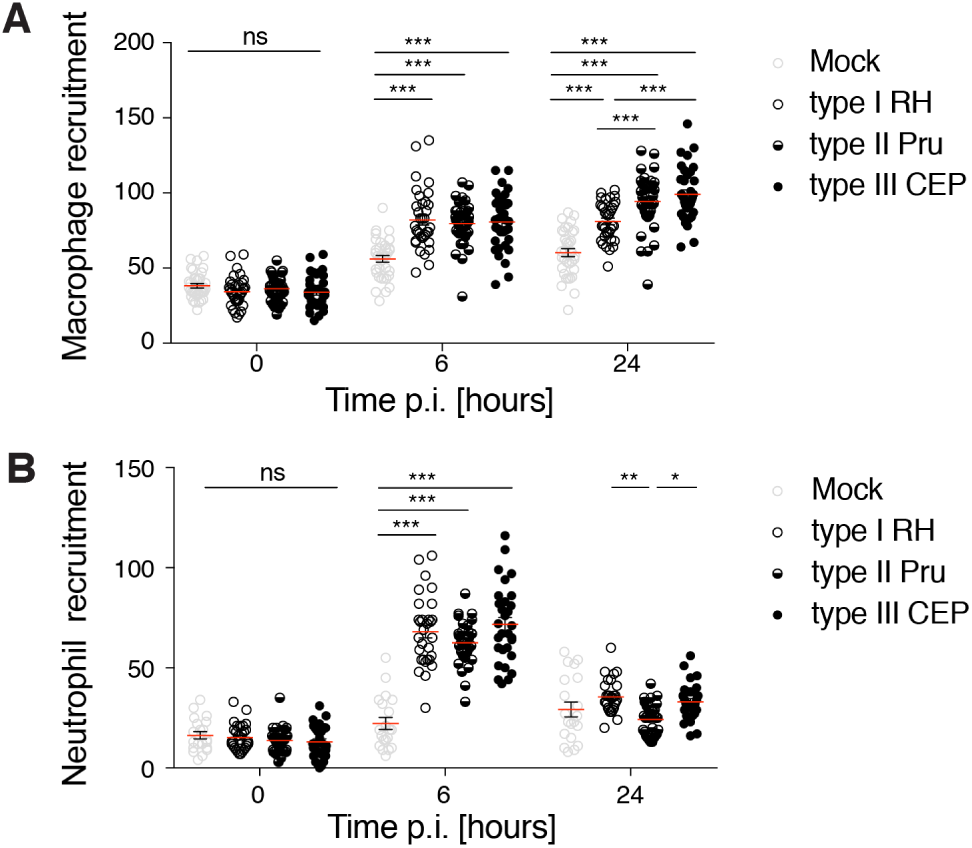
Leukocyte recruitment to *Toxoplasma gondii in vivo*. Quantification of macrophages in *mpeg1:G/U:mCherry* larvae at 0, 6 and 24 hpi injected with mock (HFF lysate, grey open circle), type I (open circle), type II (semi-closed circle) or type III (closed circle). Pooled data from at least 3 independent experiments with at least 7 larvae per condition per experiment. Mean ± SEM shown. Significance calculated using 2-way ANOVA, ns, p>0.05, ***, p *≤* 0.001. (B) Quantification of neutrophils in *lyz:dsRed* larvae at 0, 6 and 24 hpi injected with mock (HFF lysate, grey open circle), type I (open circle), type II (semi-closed circle) or type III (closed circle). Pooled data from at least 2 independent experiments with at least 3 larvae per condition per experiment. Mean ± SEM shown. Significance calculated using 2-way ANOVA, ns, p>0.05, *, p *≤* 0.01, **, p *≤* 0.01, ***, p *≤* 0.001.

To analyze *Toxoplasma*-neutrophil interactions over time, 3dpf transgenic larvae possessing red neutrophils Tg(*lyz:dsRed*)^nz50^ (herein referred to as *lyz:dsRed*), were infected with type I, II or III *Toxoplasma*-GFP, and neutrophil recruitment was quantified by fluorescent stereomicroscopy. Here, the number of neutrophils recruited to the infection site is significantly increased (~3 fold) as compared to mock injection for all three strain types at 6hpi (Fig. 3B). In contrast to macrophages, which remain at the infection site by 24hpi, the number of neutrophils recruited to the infection site (for all three strain types) is significantly decreased to basal levels by 24hpi.

### Macrophages control parasite burden *in vivo*

To analyze the interactions between type I *Toxoplasma* and macrophages in depth, we imaged infected *mpeg1:G/U:mCherry* larvae with *Toxoplasma*-GFP at 6hpi using confocal microscopy and 3D CLEM. In this case, the majority of intact type I parasites contained within macrophages are single tachyzoites inside PVs (as judged by host mitochondria association to the membrane surrounding the parasites) (Sup. Fig. 3A). This suggests that macrophages may prevent parasite replication. To follow the fate of type I parasites engulfed by macrophages in real-time, *mpeg1:G/U:mCherry* larvae infected with *Toxoplasma*-GFP were imaged by time-lapse confocal microscopy. In this case, we frequently observed the engulfment of parasites by macrophages followed by loss of GFP fluorescence, suggesting active parasite degradation (Fig. 4A, Movie 3). Consistent with this, 3D CLEM showed parasite degradation inside macrophages, as identified by fragmentation of tachyzoite organelles (Fig. 4B, Sup. Fig. 3B).

**Fig. 4.**
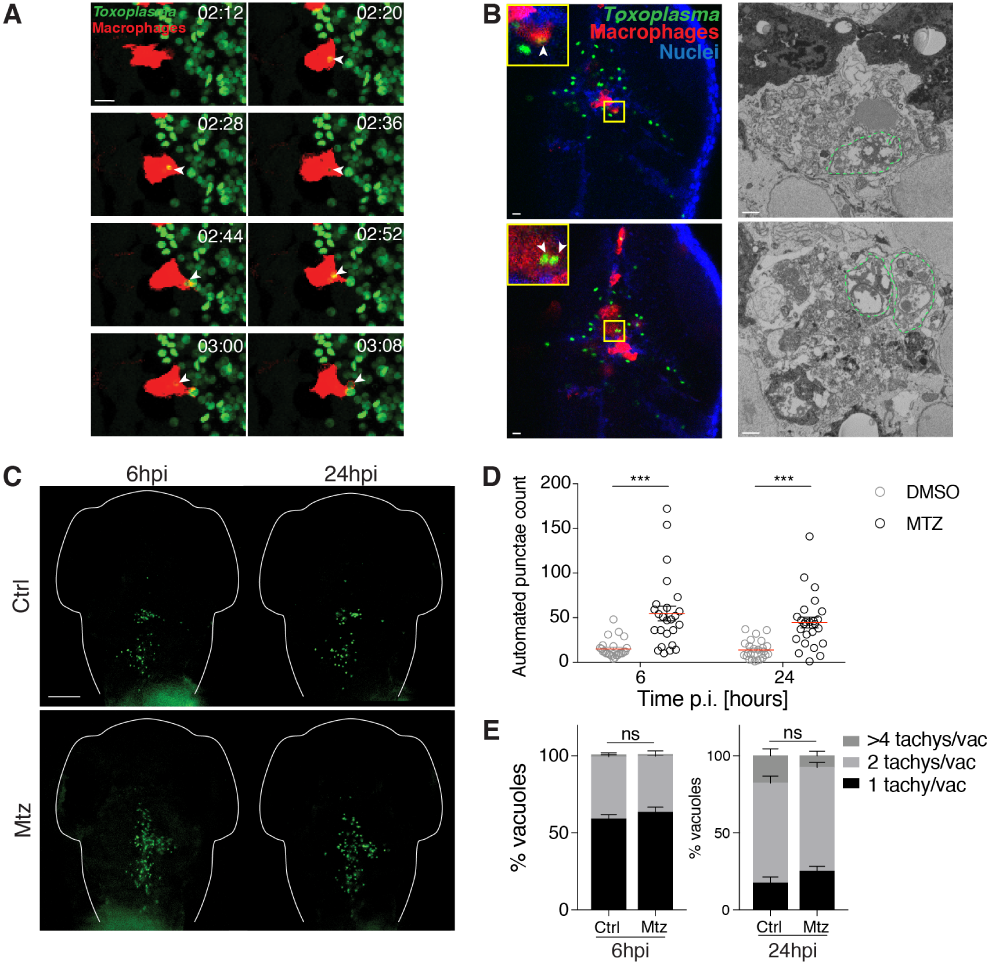
Macrophages control *Toxoplasma gondii* burden *in vivo*. (A) Representative frames extracted from *in vivo* confocal imaging of *mpeg1:G/U:mCherry* (red) larvae injected with type I *Toxoplasma*-GFP (green). First frame at 2 h 12 minutes pi followed by seven consecutive frames taken at 8-minute intervals. Showing a maximum projection of 24 z frames taken at 2 µm optical sections. Scale bar, 10 µm. See also Movie 3. (B) CLEM of dead/dying tachyzoites in the HBV of *mpeg1:G/U:mCherry* (red) larvae infected with type I *Toxoplasma*-GFP (green) at 6 hpi. Representative images extracted from confocal z-stacks of a full vibratome section (left panel) and from consecutive 50 nm SBF SEM slices of a segment of it (right panel). Dead/dying parasites indicated by white arrowheads (inset, left panels) and outlined by green dashed lines (right panels). Scale bar, 10 µm (left) and 1 µm (right). (C) Representative images of Ctrl (top panels) or macrophage-ablated (bottom panels) *mpeg:G/U:mCherry* larvae infected in HBV with type I *Toxoplasma*-GFP (green). Individual larvae were imaged and monitored at 6 and 24 hpi (from left to right) by fluorescent stereomicroscopy. Scale bar, 100 µm. (D) Automated enumeration GFP-positive punctae in the HBV at 6 and 24 hpi of Ctrl (open circle) or macrophage-ablated (closed circle) larvae infected with type I *Toxoplasma* tachyzoites. Pooled data from 3 independent experiments with at least 7 larvae per condition per experiment. Significance calculated using 2-way ANOVA, *, p *≤* 0.05, **, p *≤* 0.01. (E) Pixel volume quantification of individual GFP-positive punctae in Ctrl or macrophage-ablated larvae at 6 and 24 hpi. Presented as percentage of total vacuoles counted in the HBV that are 1 tachyzoite/vacuole (<50 pix^3^), 2 tachyzoites/vacuole (50<100 pix^3^) or >4 tachyzoites/vacuole (>100 pix^3^). Pooled data from 3 independent experiments with at least 3 larvae per time point. Significance calculated using 2-way ANOVA, ***, p *≤* 0.001. Mean ± SEM shown.

To test the role of macrophages in *Toxoplasma* infection *in vivo*, *mpeg1:G/U:mCherry* larvae were pre-treated with control (DMSO) or metronidazole (Mtz) to ablate macrophages (Sup. Fig. 3C and D). In the absence of macrophages, infected larvae showed 100% survival (Sup. Fig. 3E). However, parasite burden is significantly increased, suggesting macrophages are responsible for parasite clearance *in vivo* (Fig. 4C and D, Sup. Fig. 3F). Similar results are observed with type II and III strain infection of macrophage-ablated larvae (Sup. Fig. 3G). To analyze the viability of parasites that are cleared by macrophages, we performed vacuole volume quantification and show that DMSO and Mtz-treated larvae are comprised of equally replicating *Toxoplasma* tachyzoites (Fig. 4E). These data suggest that macrophages have a dominant role in clearing healthy viable parasites rather than supporting the parasite’s replicative niche.

## Discussion

Zebrafish infection models for studying eukaryotic parasites and other human pathogens are beginning to emerge (31, 32, 38). In this study, we establish a novel *Toxoplasma* infection model using zebrafish larvae to explore host-parasite interaction *in vivo*. We find the three main clonal lineages of *Toxoplasma* are able to invade zebrafish cells and replicate within their PV, and reveal that macrophages are key in controlling viable parasites *in vivo*.

Using confocal microscopy and CLEM, we visualize single and replicating type I tachyzoites in the zebrafish HBV exhibiting host mitochondrial association. The relatively slow replication cycle of *Toxoplasma* observed in tissue culture cells *in vitro* (>6h) is consistent with what we observe in the zebrafish HBV. GRA2 staining and CLEM of replicating tachyzoites strongly suggests PV formation and is indicative of normal type I parasite behavior as demonstrated *in vitro* using tissue culture cells and *in vivo* using other animal models (39).

The zebrafish HBV is well established to investigate host response to infection (30–32). We do not observe *Toxoplasma* dissemination from the HBV and this allows us to monitor leukocyte-parasite interactions in a localized area. Here, type II and III strains are more efficient than type I strains at maintaining a higher infectious burden. This suggests type II and III strains may be more efficient at invading non-phagocytic cell types found in the HBV and/or evading clearance by host cells. Together with CLEM evidence we conclude that *Toxoplasma* favor replication within non-phagocytic cells (such as epithelial and neuronal cells) within the HBV of zebrafish larvae.

Both live-cell imaging and CLEM showed type I parasite uptake and clearance by macrophages. In all cases of macrophage-parasite interaction captured by CLEM, macrophages retained their fluorescence during *Toxoplasma* infection and had intact nuclei and mitochondria, indicative of a healthy host cell. Our evidence obtained from time-lapse microscopy, fixed 3D CLEM and macrophage ablation highlights active parasite clearance by zebrafish macrophages *in vivo* occurs within the first 6hpi. Therefore, future work using this zebrafish infection model could uniquely explore the precise anti-parasitic mechanisms employed by macrophages during *Toxoplasma* infection.

Both live-cell imaging and CLEM showed type I parasite uptake and clearance by macrophages. In all cases of macrophage-parasite interaction captured by CLEM, macrophages retained their fluorescence during *Toxoplasma* infection and had intact nuclei and mitochondria, indicative of a healthy host cell. Our evidence obtained from time-lapse microscopy, fixed 3D CLEM and macrophage ablation highlights active parasite clearance by zebrafish macrophages *in vivo* occurs within the first 6hpi. Therefore, future work using this zebrafish infection model could uniquely explore the precise anti-parasitic mechanisms employed by macrophages during *Toxoplasma* infection.

In murine *in vivo* models, various leukocytes have been implicated in trafficking *Toxoplasma* from the site of infection. Examples include infected neutrophils that pass from the intestine to the lumen (11), as well as infected macrophages and dendritic cells that pass the blood brain barrier (40). In light of this, it is intriguing to note that our study using time-lapse microscopy shows a minority of parasite-infected macrophages moving through the brain tissue for possible parasite transport (Sup. Fig. 4A). Careful analysis of our 3D CLEM data also identified an actively replicating type I parasite inside a macrophage exhibiting host mitochondrial association to the PV (Sup. Fig. 4B-C, Movie 4). This observation is reminiscent of parasite replication “hot spots” described *in vivo* in the murine intestinal villi (11). Overall, it is remarkable that zebrafish macrophages during *Toxoplasma* infection *in vivo* have the capacity to phenocopy known behavior exhibited by murine macrophages during *Toxoplasma* infection *in vivo*. This is the case for both the type of events observed (e.g. parasite killing, trafficking, sustaining replication), and their approximate *in vivo* frequency.

In summary, we here establish a novel animal model for studying the *in vivo* innate immune response to *Toxoplasma* infection, and compare host response to the three main *Toxoplasma* strain types *in vivo*. We also discover the dominant role of macrophages in parasite clearance. Having established a zebrafish model of *Toxoplasma* infection, we have made available a unique *in vivo* infection platform for CRISPR targeting and high-throughput drug screens that, together with time-lapse microscopy, can be used to identify determinants underlying *Toxoplasma* infection control.

## Supporting information

Sup Movie1. In vivo replication of Toxoplasma gondii

Sup Movie 2. 3D CLEM of Toxoplasma gondii replication in the zebrafish hindbrain

Sup Movie 3. Phagocytosis of Toxoplasma gondii by a macrophage

Sup Movie 4. 3D CLEM of replicative Toxoplasma gondii inside a zebrafish macrophage

Sup Movie 5. 3D visualization of GFP-positive vacuole in the zebrafish hindbrain by confocal microscopy for volume pixel analysis

## ACKNOWLEDGEMENTS

We thank Vincenzo Torraca, Gina Duggan, Margarida Castro Gomes, Joseph Wright and Barbara Clough for scientific discussions. We thank Moritz Treeck for providing the α-GRA2 antibody.

## Author Contributions

NY conducted all experiments with the exception of the electron microscopy, MCD conducted all sample preparation, imaging and image analysis for electron microscopy, MJM initiated the infection model, LC provided electron microscopy expertise and equipment, EMF and SM supervised the study, NY, MCD, EMF and SM wrote the manuscript and all authors provided input.

## Materials and Methods

### Ethics statement

Animal experiments were performed according to the Animals (Scientific Procedures) Act 1986 and approved by the Home Office (Project licenses: PPL P84A89400 and P4E664E3C). All experiments were conducted up to 5 days post-fertilization.

#### Zebrafish husbandry and maintenance

Fish were reared and maintained at 28.5°C on a 14hr light, 10hr dark cycle. Embryos obtained by natural spawning were maintained in 0.5x E2 media supplemented with 0.3 µg/ml methylene blue. Larvae were anesthetized with 20 µg/ml tricaine (Sigma-Aldrich) during the injection procedures and for live *in vivo* imaging. All experiments were carried out on TraNac background (41) larvae to minimize obstruction of fluorescence signal by pigmentation leading to misrepresentation in parasite dose quantification.

#### Parasite culture, preparation and infection

*Toxoplasma* (RH/Pru/CEP) expressing GFP/luciferase or Tomato was maintained *in vitro* by serial passage on human foreskin fibroblasts (HFFs) cultures (ATCC). Cultures were grown in DMEM high glucose (Life Technologies) supplemented with 10% FBS (Life Technologies) at 37°C in 5% CO_2_. Parasites were prepared from 25G followed by 27G syringe-lysed HFF cultures in 10% FBS. Excess HFF material removed by centrifugation for 10min at 50 x g. After washing with PBS, *Toxoplasma* tachyzoites were resuspended at 2×10^6^ tachyzoites/µl in PBS. During injection, tachyzoites were maintained at room temperature and passed through 29G myjector syringe (Terumo) to dissociate clumps and homogenize the suspension. Control infections were carried out using uninfected HFF cultures prepared as described above. 3dpf larvae were anesthetized and injected with ~2.5nl of parasite suspension into the HBV. HBV injections were carried out as previously described (42). Larvae are optimally maintained at 28.5°C, but develop normally between 23-33°C (43). *Toxoplasma* invades and replicates at a minimum of 33°C. Infected larvae were therefore transferred into media pre-warmed to 33°C to ensure normal zebrafish development and parasite replication (Sup. Fig. 1A). Progress of infection was monitored by fluorescent stereomicroscopy (Leica M205FA, Leica Microsystems).

#### Quantification of parasite dose and burden

For parasite dose quantification z-stack images of the infected hindbrain were taken within 5-10min using the Leica M205FA fluorescent stereomicroscope on a 130x magnification using a 1x objective. Images were analyzed using the particle analysis function in Fiji software (44). For manual quantification of parasite burden, z-stack images were taken using the Leica M205FA. GFP-positive punctae were quantified using the multi-point tool in Fiji software. Computer vision driven automated parasite burden quantifications were carried out using the ZedMate plugin in Fiji software to corroborate manual quantifications (37). Pixel volume quantifications were carried out by 3D projecting confocal z-stack images and using the 3D objects counter tool in Fiji (Movie 5).

#### Live imaging, image processing and analysis

Live *in vivo* imaging was performed on anesthetized larvae immobilized in 1% low melting-point agarose in 35mm glass-bottomed dishes (MatTek Corp.). Widefield microscopy was performed using a 40x objective. Z-stacks were acquired at 10-minute intervals. 60 z slices were taken at 2µm sections. Confocal microscopy was performed using the Zeiss Invert LSM 710 (Carl Zeiss AG) and the LSM 880 (Carl Zeiss AG) using a 40x and 63x objective. Z-stacks were acquired at 8-minute intervals. 60 z slices were taken at 0.9µm sections per larva. For all time-lapse acquisitions, larvae were maintained at 33°C. For mitochondria staining, larvae were injected with 1nl MitoTracker® DeepRed (250µM, Life Technologies) 40min prior to embedding for live confocal microscopy.

#### Wholemount immunohistochemistry

Euthanized larvae were fixed overnight at 4°C in 4% paraformaldehyde supplemented with 0.4% Triton X-100 and washed in PBS, 0.4% Triton X-100 before staining. Briefly, after a 20min wash in PBS 1% Triton X-100, larvae were incubated overnight at 4°C in blocking solution: PBS, supplemented with 10% FBS, 1% DMSO and 0.1% Tween 20. Primary antibodies diluted in blocking solution were applied overnight at 4°C. Larvae were washed 4×15min with PBS supplemented with 0.1% Tween 20. Secondary antibodies diluted in blocking solution were applied overnight at 4°C. Larvae were washed 4×15min with PBS supplemented with 0.1% Tween 20. Hoechst 33342 staining of larvae was carried out at room temperature for 10min, followed by 3×10min washes with PBS, 0.1% Tween 20. Larvae were then cleared through a glycerol series before imaging by confocal microscopy (Zeiss LSM 710).

#### Antibodies

Primary antibodies used were mouse α-GRA2 (BIO.018.5, BIOTEM) kind gift of Moritz Treeck, The Francis Crick Institute, UK. Secondary antibodies used were goat α-mouse AF647 (Invitrogen).

#### 3D correlative light and electron microscopy (CLEM)

For serial blockface scanning electron microscopy (SBF SEM), euthanized larvae were fixed overnight at 4°C in 4% formaldehyde (Taab Laboratories Equipment Ltd.). Hoechst 33342 staining of larvae was carried out at room temperature for 10min without permeabilization and larvae were subsequently washed 3×10min with 0.1M phosphate buffer (PB). Larvae were embedded in 3% low-melt agarose in 35mm glass-bottomed dishes. Larvae were covered in 0.1M PB for high-resolution confocal microscopy (Zeiss LSM 710). Larvae were maintained in 1% formaldehyde in 0.1M PB until further processing. The embedded larvae were sectioned using a Leica VT1000 S vibrating blade microtome (Leica Biosystems). 50µm sections were collected and stored in 0.1M PB in a 24-well glass-bottomed plate (MatTek Corp.). The sections were imaged again using a Zeiss Invert 710 LSM confocal (Carl Zeiss AG) and a 20x Ph2 objective. The sections containing *Toxoplasma* were then processed following the method of the National Centre for Microscopy and Imaging Research (45). In brief they were post-fixed in 2.5% (v/v) gluteraldehyde/4% (v/v) formaldehyde in 0.1M PB for 30min at room temperature, stained in 2% osmium tetroxide/1.5% potassium ferricyanide for 1h on ice, incubated in 1% w/v thiocarbohydrazide for 20min before a second staining with 2% osmium tetroxide, and incubation overnight in 1% aqueous uranyl acetate at 4°C. Sections were stained with Walton’s lead aspartate for 30min at 60°C and dehydrated stepwise through an ethanol series on ice, incubated in a 1:1 propylene oxide/Durcupan resin mixture and embedded in Durcupan ACM® resin according to the manufacturer’s instructions (Sigma-Aldrich). Blocks were trimmed to a small trapezoid, excised from the resin block and attached to a SBF SEM specimen holder using conductive epoxy resin (Circuitworks CW2400). Prior to commencement of a SBF SEM imaging run, the sample was coated with a 2nm layer of platinum to further enhance conductivity using a Q150R S sputter coater (Quorun Technologies Ltd.).

SBF SEM data was collected using a 3View2XP (Gatan Inc.) attached to a Sigma VP SEM (Zeiss). Inverted backscattered electron images were acquired through the entire extent of the region of interest. For each 50nm slice, a low-resolution overview image (pixel size of 50nm using a 1.5µs dwell time) and several high-resolution images of the different regions of interest (indicated magnification 5000x, pixel size of 6-7nm using a 1.5µs dwell time) were acquired. The overview image was used to relocate the region of interest defined by the confocal images of the sections. The SEM was operated in variable pressure mode at 5Pa. The 30µm aperture was used, at an accelerating voltage of 2kV. Typically, between 300 and 1000 slices were necessary for an entire region of interest.

As data was collected in variable pressure mode, only minor adjustments in image alignment were needed, particularly where the field of view was altered in order to track the cell of interest. All the images were converted to tiff in Digital Micrograph (Gatan Inc.), and tiff stacks were automatically aligned using TrakEM2, a FIJI framework plug-in (Cardona et al. 2012). Manual segmentations were done in TrakEM2. For Fig. 1D, labels were exported as tiff for visualization in 3D in ClearVolume, a FIJI framework plug-in (46). For Sup. Fig. 4B, they were exported as Amira labels for visualization in 3D in Amira Software (Thermo Fisher Scientific). Movie 2 was generated in FIJI, Movie 4 in Amira, both were compressed in Quick Time Pro with the H.264 encoder.

#### Measurement of leukocyte recruitment to the site of infection

Anesthetized larvae were imaged 0, 6 and 24hpi by fluorescent stereomicroscopy (Leica M205FA). 15 z slices were taken at 130x magnification. Images were further analyzed using Fiji software.

#### Metronidazole targeted macrophage depletion

Dechorionated 2dpf TraNac-Tg(*mpeg1:Gal4-FF*)^*gl25*^/Tg(*UAS-E1b:nfsB.mCherry*)^*c264*^ larvae were placed in embryo media supplemented with metronidazole (10mM, Sigma-Aldrich), 1% DMSO. Larvae were then placed in fresh 10mM metronidazole solution at 33°C post-infection. Control-treated larvae were maintained in embryo media supplemented with 1% DMSO.

#### Statistical analysis

Significance testing was performed by Student t-test, 1-way ANOVA or by 2-way ANOVA. The level of significance is shown as ns, p > 0.05; *, p *≤* 0.05; **, p *≤* 0.01; ***, p *≤* 0.001.

## Appendix 1. Supplementary Information

This is a supplementary section to the preprint manuscript by Yoshida *et al.* 2019.

**Supplementary Figure 1.**
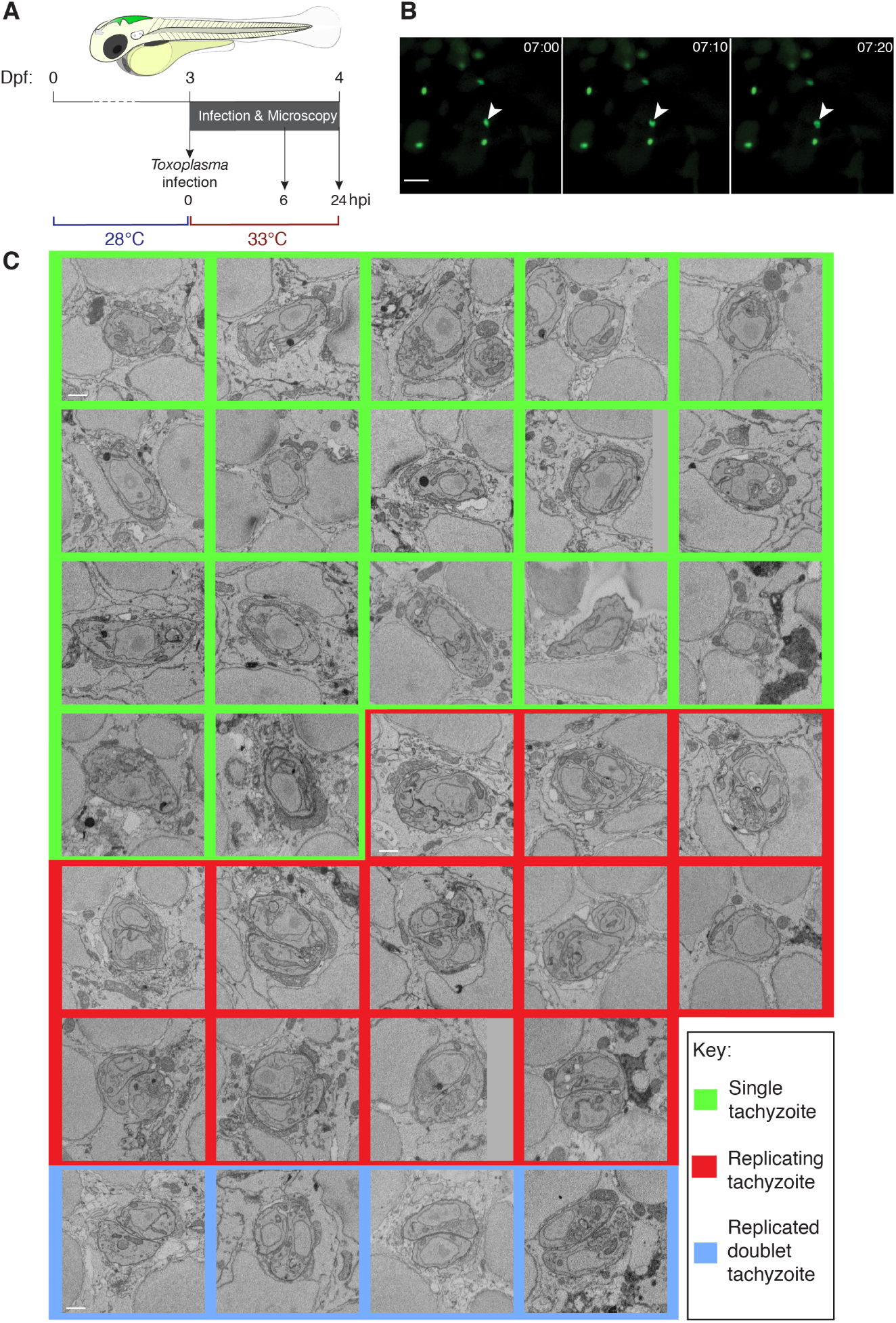
*Toxoplasma gondii* tachyzoites are intracellular and replicate in zebrafish. (A) Schematic of the infection model utilized with a cartoon of the zebrafish larva (3dpf) showing the site of *Toxoplasma* tachyzoite (green) injection in the hindbrain ventricle (HBV). Infected larvae were maintained at 33°C post-injection and monitored up to 24 hours post-infection (hpi). (B) Representative frames extracted from *in vivo* widefield imaging of larvae injected with type I *Toxoplasma*-GFP (green). First frame at 7hpi followed by two consecutive frames taken at 10-minute intervals. Showing a single z plane from 60 taken at 2 µm optical sections. Scale bar, 20 µm. See also Movie 1. (C) CLEM of tachyzoites in the HBV of *mpeg1:G:U:mCherry* larvae infected with type I *Toxoplasma*-GFP at 6hpi. Representative images of 33/36 *Toxoplasma* in zebrafish host cells extracted from consecutive 50 nm SBF SEM slices. *Toxoplasma* tachyzoites were imaged in their full volume to accurately determine their stage. Single (green box), replicating (red box) and replicated doublet (blue box) tachyzoites were observed. Scale bar, 1 µm.

**Supplementary Figure 2.**
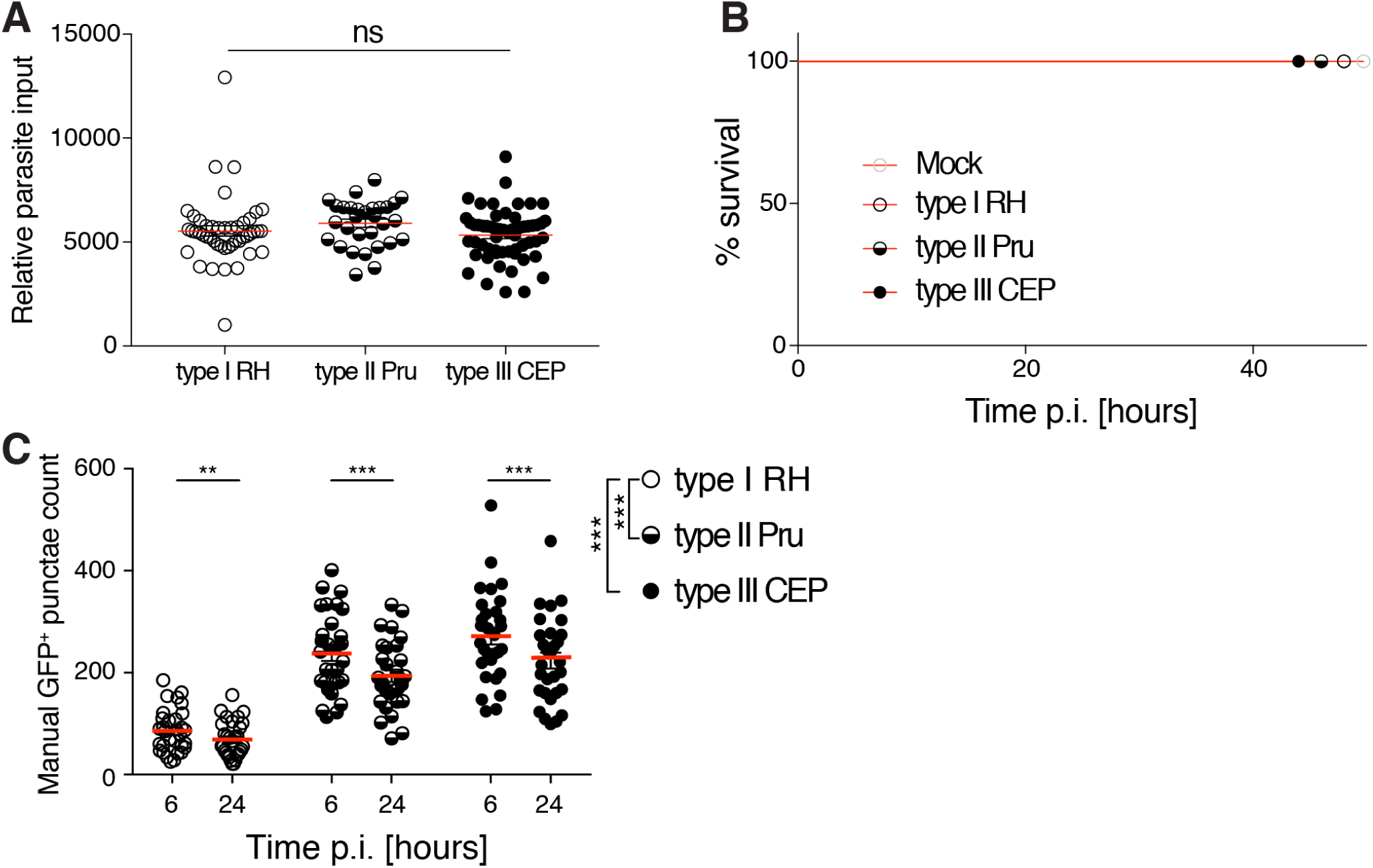
The zebrafish larvae model of acute *Toxoplasma gondii* infection is non-lethal. (A) Quantification of type I (open circle), type II (semi-closed circle) or type III (closed circle) dose of *Toxoplasma*-GFP using particle analysis of infected HBV images obtained by fluorescent stereomicroscopy at 0 hpi. Mean ± SEM shown. Significance calculated using Student’s t-test, ns, p>0.05. (B) Survival curves of larvae injected with mock (human foreskin fibroblast, HFF, lysate), type I, type II or type III *Toxoplasma* tachyzoites. Pooled data from at least 3 independent experiments with at least 5 larvae per condition per experiment. (C) Manual enumeration of GFP-positive punctae in the HBV at 6 and 24 hpi of type I (open circle), type II (semi-closed circle) or type III (closed circle) *Toxoplasma*-GFP. Mean ± SEM shown. Pooled data from at least 3 independent experiments with at least 5 larvae per condition per experiment. Significance calculated using 2-way ANOVA, **, p *≤* 0.01, ***, p *≤* 0.001.

**Supplementary Figure 3.**
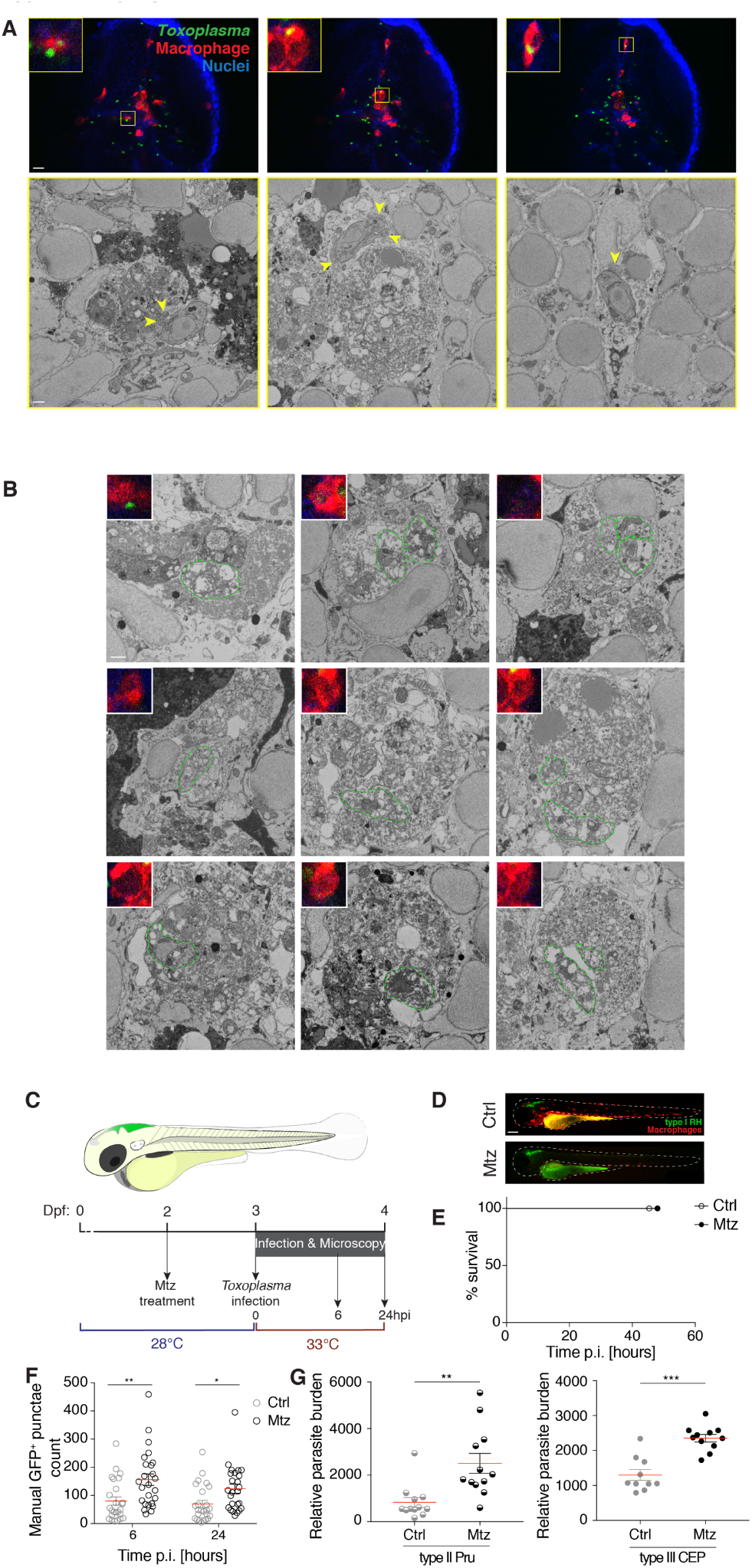
Zebrafish macrophages control *Toxoplasma gondii*. (A) CLEM of parasites inside macrophages in the HBV of mpeg1:G/U:mCherry (red) larvae infected with type I *Toxoplasma*-GFP (green) at 6hpi. Representative images extracted from 44 confocal z-stacks of a full section (top panels). The localizations of respective high-resolution SBF SEM images (bottom panels) are denoted with yellow boxes. Host mitochondrial recruitment to the parasitophorous vacuole indicated by yellow arrowheads. Scale bars, 20 µm (top panels) and 1 µm (bottom panels). (B) CLEM of putative dead tachyzoites in the HBV of mpeg1:G/U:mCherry (red) larvae infected with type I *Toxoplasma*-GFP (green) at 6hpi. Representative images extracted from confocal z-stacks of a full section (inset) and from consecutive 50 nm SBF SEM slices of a segment of it. Only the first parasite (top left) showed GFP fluorescence. Putative dead parasites indicated by green dashed outline. Scale bar, 1 µm. (C) Schematic of the infection model utilized with a cartoon of the zebrafish larva (3 dpf) showing the site of *Toxoplasma* tachyzoite (green) injection in the HBV. Prior to infection, larvae were pre-treated from 2 dpf with DMSO (Ctrl) or metronidazole (Mtz). Infected larvae were maintained at 33°C post-infection and monitored up to 24hpi. (D) Representative images of Ctrl or Mtz treated *mpeg1:G/U:mCherry* larvae (red) injected with type I *Toxoplasma*-GFP (green) 0 hpi. Scale bar, 200 µm. (E) Survival curves of Ctrl (open circles) or macrophage-ablated (metronidazole treated *mpeg1G/U:mCherry*, closed circles) larvae infected in the HBV with type I *Toxoplasma*-GFP. Pooled data from 3 independent experiments with at least 7 larvae per condition per experiment. (F) Manual enumeration GFP-positive punctae in the HBV at 6 and 24hpi of Ctrl (grey) or macrophage-ablated (black) larvae infected with type I *Toxoplasma*-GFP. Mean ± SEM shown. Pooled data from 3 independent experiments with at least 7 larvae per condition per experiment. Significance calculated using 2-way ANOVA, *, p ≤ 0.05, **, p ≤ 0.01. (G) Quantification of relative parasite burden in the HBV of Ctrl (grey) or macrophage-ablated (black) larvae infected with type II (semi-closed circle) or III (closed circle) 24hpi. Mean ± SEM shown. Showing 1 representative experiment of 3 with at least 10 larvae per condition per experiment. Significance calculated using Student’s t-test, **, p *≤* 0.01, ***, p *≤* 0.001.

**Supplementary Figure 4.**
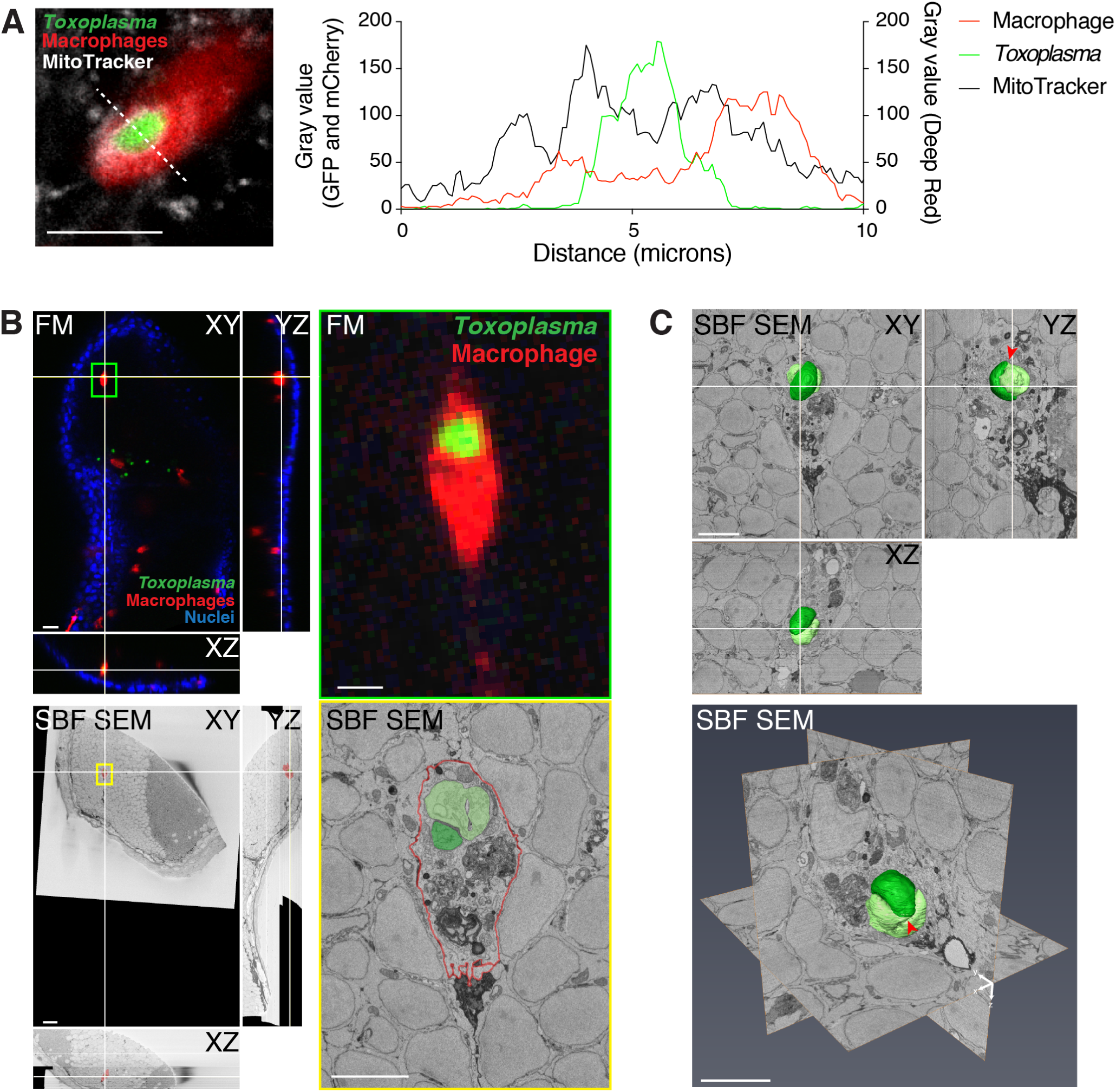
Replicative *Toxoplasma gondii* inside a zebrafish macrophage. (A) CLEM of parasite replication inside a macrophage in the HBV of *mpeg1:G/U:mCherry* (red) larvae infected with type I *Toxoplasma*-GFP (high dose, green) at 6hpi. Orthoslices of 44 confocal z-stacks of a full vibratome section (FM, left top panel) and of 1662 consecutive 50 nm SBF SEM slices of a segment of it (left bottom panel). Color boxes show localization of the confocal (right top, green) and high-resolution SBF SEM images (right bottom, yellow). The replicating *Toxoplasma* (green and light green) and plasma membrane of the macrophage (red) were manually segmented. Blue arrowhead in the light green *Toxoplasma* indicate 2 nuclei. Host mitochondrial recruitment to the parasitophorous vacuole indicated by yellow arrowheads. Scale bars, 20 µm (left) and 5 µm (right). (B) Orthoslices (top panel) and 3D view (bottom panel) of the 3D model of the 2 segmented type I *Toxoplasma* shown in (A) overlaid on 373 consecutive 50 nm SBF SEM slices. Area where the 2 *Toxoplasma* are still joined indicated by the red arrowhead, nucleus of the macrophage by the blue arrow, host mitochondrial recruitment to the parasitophorous vacuole by yellow arrowheads. Scale bars, 5 µm. See also Movie 4. (C) Representative image extracted from a z-stack from confocal imaging of live *mpeg1G/U:mCherry* larvae infected in the HBV with type I *Toxoplasma*-GFP and stained with MitoTracker at 6hpi and the fluorescent intensity profile of a parasite exhibiting host mitochondrial association within a macrophage. Scale bar, 10 µm.

**Movie 1. *In vivo* replication of *Toxoplasma gondii*.** *In vivo* fluorescent widefield imaging of larvae injected with type I *Toxoplasma*-GFP (green). First frame at 3h 50 minutes post-infection (mpi) followed by frames taken at 10-minute intervals until 8h 30mpi. Showing a single z plane from 60 taken at 2 µm optical sections. Scale bar, 20 µm.

**Movie 2. 3D CLEM of *Toxoplasma gondii* replication in the zebrafish hindbrain.** SBF SEM of tachyzoites in the HBV of larvae injected with type I *Toxoplasma*-GFP at 6hpi. Representative examples of single (left), replicating (middle) and replicated doublet (right) *Toxoplasma* from a total of 36 found in zebrafish cells (see Fig. 1D and Sup. Fig. 1C) imaged in their whole volume to accurately determine their stage. 112 consecutive 50 nm SBF SEM slices of a different segment of a section for each example. Scale bar, 1 µm.

**Movie 3. Phagocytosis of *Toxoplasma gondii* by a macrophage.** *In vivo* confocal imaging of mpeg1:G/U:mCherry larvae injected with type I *Toxoplasma*-GFP (green). First frame at 1h 48mpi followed by frames taken at 8-minute intervals until 3h 48mpi. Showing a maximum projection of 24 z frames from 60 taken at 2 µm optical sections. Scale bar, 10 µm.

**Movie 4. 3D CLEM of replicative *Toxoplasma gondii* inside a zebrafish macrophage.** CLEM of intraphagocytic parasite replication in the HBV of *mpeg1:G/U:mCherry* larvae infected with type I *Toxoplasma*-GFP at 6hpi. 373 consecutive 50 nm SBF SEM slices on which the replicating *Toxoplasma* (green and light green) and plasma membrane of the macrophage (red) were manually segmented. A surface was generated to build a 3D model of the 2 segmented *Toxoplasma* in Amira Software. Scale bar, as indicated.

**Movie 5. 3D visualization of GFP-positive vacuole in the zebrafish hindbrain by confocal microscopy for volume pixel analysis.** Showing representative 3D confocal images of GFP-positive vacuoles (green) in fixed larvae infected with type I *Toxoplasma*-GFP from 6 to 24hpi. Volumes were categorized into 1 tachyzoite/vacuole (<50 pix^3^), 2 tachyzoites/vacuole (50<100 pix^3^) or >4 tachyzoites/vacuole (>100 pix^3^). Vacuole volume measured is indicated by the yellow outline. Scale bar, 20 µm.

